# Inhibition of *Slc17a7* expressing neurons in the basolateral amygdala which project to the nucleus accumbens shapes the fidelity of motivated behavior

**DOI:** 10.64898/2026.01.26.701876

**Authors:** William D. Mercer, Iltan Aklan, Nathaniel E. Connolly, Shivangi M. Inamdar, Benjamin L. Fisher, Chase M. Larsson, Kyle H. Flippo

## Abstract

Adaptive behavior depends on the ability to form predictive associations between environmental cues and biologically relevant outcomes. Classical models posit that projections from the basolateral amygdala (BLA) to the nucleus accumbens (NAc) promote reward seeking, whereas projections to the central amygdala facilitate defensive responding. However, accumulating evidence indicates that molecular identity and inhibitory dynamics introduce additional layers of functional heterogeneity within these pathways. Genetically defined subsets of BLA→NAc neurons can drive either reward-related or aversive behaviors, and single-cell imaging studies show that cue-responsive BLA ensembles undergo an inhibitory shift as conditioning progresses. How these molecular and inhibitory mechanisms interact within defined projections to shape motivated behavior remains unclear. To address this, we examined Vglut1-expressing BLA→NAc neurons (Vglut1^BLA→NAc^), the predominant excitatory population within this pathway. Whole-cell electrophysiology revealed that reward conditioning increases inhibitory synaptic input onto Vglut1^BLA→NAc^ neurons and reduces intrinsic excitability. Using a dual-recombinase chemogenetic strategy, we found that selective inhibition of this projection enhances acquisition of a cued reward task, increases instrumental reward seeking, and elevates reward valuation in a two-choice preference assay. Similar enhancement was observed in a fear-discrimination task, where inhibition of Vglut1^BLA→NAc^ neurons improved differentiation between threat- and safety-associated cues. These findings identify inhibitory regulation of Vglut1^BLA→NAc^ neurons as a key mechanism that strengthens stimulus–outcome associations. By demonstrating that conditioning-associated inhibitory plasticity modulates this major BLA output pathway, the work refines prevailing models of BLA→NAc function and highlights inhibition as an important contributor to the fidelity of motivated behavior.

## 1. Introduction

The ability to use environmental stimuli as predictors of food availability or potential threats is essential for survival. Formation of such associations relies on neural mechanisms that enable organisms to form predictive links between stimuli and outcomes ^[1, 2]^. A central component of these processes involves coordinated interactions between limbic and striatal systems, particularly projections from the basolateral amygdala (BLA) to the nucleus accumbens (NAc). The BLA integrates sensory and affective information and, through projections to the NAc (BLA→NAc), guides motivated behavior^[3-5]^. Classical models have emphasized that anatomically defined BLA projection pathways determine their functional roles in associative learning. In this framework, BLA→NAc neurons promote reward seeking and approach behavior, consistent with findings that excitation of this pathway enhances motivation and conditioned responding to reward-predictive cues or actions^[6-12]^. In contrast, BLA→CeA neurons are proposed to facilitate defensive behaviors and enhance fear learning^[7]^ and have long-been thought to be required for Pavlovian fear conditioning^[13-15]^. These projection-specific functions have been further interpreted as encoding the predictive valence of environmental stimuli and associated outcomes^[16-20]^, spanning reward to threat, to optimize behavioral decisions. A substantial body of work supports the core features of this projection-based organizational model.

More recent evidence, however, suggests that molecular identity introduces an additional layer of functional specialization within defined projections^[21, 22]^. Studies examining genetically defined subsets of BLA→NAc neurons demonstrate that molecularly distinct populations can exert opposing influences on behavior. For example, activation of *Rspo2*-expressing BLA→NAc neurons promotes defensive responses and suppresses reward conditioning^[21]^, with similar findings reported for the overlapping *Fezf2*-expressing population^[22]^. These results indicate that BLA→NAc projections are not functionally uniform but instead comprise heterogeneous subpopulations whose molecular profile shapes their contribution to motivated behavior. This heterogeneity further implies that the balance of excitation and inhibition across these subpopulations is likely critical for selecting appropriate behavioral responses across diverse environmental contingencies.

Complementing this molecular perspective, single-cell in vivo calcium imaging studies reveal that BLA population dynamics shift systematically during behavioral conditioning. As animals acquire reward–predictive associations, an increasing proportion of BLA neurons become inhibited by reward-predictive cues, whereas cue-evoked excitation remains relatively stable^[23, 24]^. This inhibitory shift is stimulus-specific and recruits previously unresponsive neurons into an “inhibitory ensemble,” suggesting that dynamic inhibitory modulation is a hallmark of later conditioning stages once task performance stabilizes. Together, these findings point to an emerging view in which both molecular identity and conditioning-related inhibitory dynamics shape BLA function during motivated behavior.

Despite this growing evidence, how inhibitory signaling within molecularly defined BLA→NAc projections contributes to motivated behavior remains poorly understood. To address this question, we performed whole-cell electrophysiological recordings in Vglut1 (*Slc17a7*)-expressing BLA→NAc neurons (Vglut1^BLA→NAc^), the predominant excitatory population comprising this pathway (∼70% of BLA→NAc neurons^[25]^). We tested whether reward conditioning alters intrinsic excitability and inhibitory synaptic inputs onto these neurons. Reward conditioning increased inhibitory input and reduced excitability in Vglut1^BLA→NAc^ neurons. To determine the behavioral significance of this adaptation, we selectively inhibited this projection using a dual recombinase chemogenetic strategy. Inhibition of Vglut1^BLA→NAc^ neurons markedly enhanced acquisition of a cued reward task, increased instrumental reward seeking, and elevated reward valuation in a two-choice preference assay. Parallel behavioral enhancement was observed in a fear-discrimination paradigm, where inhibition of this projection improved differentiation between shock-predictive and safety cues. Together, these findings reveal a previously unrecognized role for inhibitory regulation within the BLA→NAc pathway in strengthening stimulus–outcome associations. By demonstrating that conditioning-associated increases in inhibitory signaling modulate the dominant excitatory projection from the BLA, this work refines prevailing models of BLA→NAc function and highlights inhibitory dynamics as a key mechanism for shaping the fidelity of motivated behavior.

## 2. Materials and Methods

### Key resources table

| Reagent type<br>or resource | Source | Identifier |
| --- | --- | --- |
| **<u>Viruses</u>** |  |  |
| pAAV-hSyn-DIO-EGFP | Addgene | Catalog: 50457- AAVrg |
| AAV pEf1a-DIO-FLPo-WPRE-hGHpA | Addgene | Catalog: 87306- AAVrg |
| pAAV-Ef1a-fDIO-mCherry | Addgene | Catalog: 114471- AAV9 |
| pAAV-hSyn-fDIO-hM4D(Gi)-<br>mCherry-WPREpA | Addgene | Catalog: 154867- AAV9 |
| **<u>Chemicals</u>** |  |  |
| DREADD Agonist 21 (C21) | Tocris Bioscience | Catalog: 5548 |
| Sucrose | Research Productions<br>International | CAS No. 2250025-92-2 |
| Paraformaldehyde (4%) | Thermoscientific | Catalog: J19943.K2 |
| Vectashield Antifade Mounting<br>Media with DAPI | (Vector Laboratories) |  |
| **<u>Antibodies</u>** |  |  |
| Donkey Anti-Rabbit IgG (H+L) Highly Cross-adsorbed Secondary Antibody, Alexa Fluor™ 555 | Invitrogen | Catalog: A-31572 |
| mCherry Polyclonal Antibody | Invitrogen | Catalog: PA5-34974 |
| **<u>Equipment</u>** |  |  |
| PyControl- Mouse Behavior Box | LabMaker |  |
| NIR Video Fear Condition (VFC) System for Optogenetics | Med Associates Inc. |  |
| Olympus Slideview VS200 | Olympus |  |
| **<u>Software</u>** |  |  |
| Pycontrol GUI | Akam, et al., 2022 | Pycontrol v2.02 |
| VideoFreeze™ Video Fear Conditioning Software | Med Associates Inc. |  |
| GraphPad Prism | Dotmatics |  |
| Olympus OlyVIA | Olympus |  |

### Animals

Vglut1-IRES2-Cre-D were obtained through Jackson Laboratories (JAX: 037512). All experiments presented in this study were conducted according to the animal research guidelines from and approved by the Iowa City Department of Veterans Affairs IACUC. Mouse were kept housed at room temperature (20-22° C) with a 12-hour light/dark cycle with standard mouse chow and water provided ad libitum, unless otherwise stated. Littermates were randomly assigned to experimental groups. For all studies we used male and female mice 10-16 weeks of age. All the animals used in this study were singly housed for at least two weeks before behavioral experiments. Any animals exhibiting abnormal health status were excluded from the study.

### Surgical Protocols

All surgical procedures were performed utilizing aseptic techniques, with mice anesthetized utilizing isoflurane (2-4%) or intraperitoneal injection of ketamine hydrochloride (100 mg/kg) and xylazine (20 mg/kg) mixture diluted in sterile saline and placed in a stereotaxic frame (Stoelting). In case of ketamine usage, atipamezole (8 mg/kg) was used to reverse effects after surgery. Subcutaneous injection of flunazine (10 mg/kg) was provided as postoperative care. Craniotomies were made above the injection sites with specific coordinates for each region as follows; NAc (AP +1.145 mm; ML ±1.25mm; DV −4.55 mm from bregma) and BLA (AP −1.38 mm; ML ±3.15mm; DV −4.65 mm from bregma). Hamilton Neurosyringes (1 µl) were used to inject 500 nL of AAV at each injection site at a rate of 50 nL/min. All viruses from Addgene were diluted to a 1:6 working solution before injection. After injection, the syringe remained at the site for a minimum of five minutes to allow for absorption into the tissue.

### Electrophysiology Recordings

For electrophysiological recordings, a cre-dependent retrogradely transported AAVrg-hSyn-DIO-eGFP was injected into the NAc of Vglut1-Cre mice (∼8 weeks old), and GFP labeled BLA neurons were recorded from. We adapted a protocol for slice preparation (Ting et al., 2018^[26]^). Sacrificed mice brains were submerged in a cold NMDG-HEPES aCSF cutting solution (in mM): 92 NMDG, 2.5 KCl, 1.25 NaH_2_PO_4_, 30 NaHCO_3_, 20 HEPES, 25 Glucose, 5 Na-Ascorbate, 2 Thiourea, 3 Na-Pyruvate, 10 MgSO_4_·7H_2_O, and 0.5 CaCl_2_·2H_2_O (pH 7.3-7.4 with HCl; 300-310 mOsm). 300 μm thick-brain slices from BLA region were obtained by using vibratome in 95% O_2_ / 5% CO_2_ aerated chilled cutting solution. Every slice was transferred to the NMDG-HEPES aCSF solution at 37 °C, and 50 µM NaCl solution was gradually dropped into the aCSF over 5 min. A couple of minutes after the last NaCl addition, BLA slices were transferred to a 95% O_2_ / 5% CO_2_ aerated holding solution at 25 °C containing (in mM): 92 NaCl, 2.5 KCl, 1.2 NaH_2_PO_4_, 30 NaHCO_3_, 20 HEPES, 25 Glucose, 5 Na-Ascorbate, 2 Thiourea, 3 Na-Pyruvate, 2 MgSO_4_·7H_2_O, and 2 CaCl_2_·2H_2_O (pH 7.3-7.4; 300-310 mOsm). After the incubation period (at least 45 minutes), brain sections were placed in a recording chamber perfused with an aCSF solution containing (in mM): 124 NaCl, 2.5 KCl, 1.2 NaH_2_PO_4_, 24 NaHCO_3_, 5 HEPES, 12.5 Glucose, 2 MgSO_4_·7H_2_O, and 2 CaCl_2_·2H_2_O (pH 7.3-7.4; 300-310 mOsm).

Loose seal and whole cell recordings were obtained from NAc-projecting BLA neurons in naive and conditioned Vglut1-Cre mice. In voltage clamp mode of whole recordings, spontaneous inhibitory post synaptic currents (sIPSC) were recorded under 10 µM CNQX and 50 µM AP5 (no PTX) with a CsCl internal solution containing (in mM): 125 CsCl, 5 NaCl, 10 HEPES, 0.6 EGTA, 10 Lidocaine Ethyl Bromide (QX-316), 4 Mg-ATP, and 0.4 Na_2_-GTP (pH with KOH to 7.3-7.4; 285-295 mOsm). In current clamp mode, whole cell recordings were obtained under synaptic blockers (10 µM CNQX, 50 µM AP5, and 50 µM PTX) with a K^+^ Gluconate internal solution containing (in mM): 130 K^+^ D-Gluconate, 4 KCl, 10 HEPES, 0.3 EGTA, 10 Na_2_Phosphocreatine, 4 Mg-ATP, and 0.4 Na2-GTP (pH with KOH to 7.3-7.4; 285-295 mOsm). From these recordings, resting membrane potential after immediately breaking into the cell (Vm) was obtained. Currents were injected in 30 pA steps (0 to 120 pA), and 1^st^ spike latency and number of action potentials were determined.

Analysis and data collection were performed with MultiClamp 700B Amplifier (Molecular Devices, San Jose, CA) and Axon pCLAMP 11.2.2 software (Molecular Devices, San Jose, CA).

### Immunofluorescence

For immunohistochemistry brains were collected and fixed for 24 h in 4% paraformaldehyde (PFA) and transferred to 30% sucrose for at least 48 h and until sinking. Brains were cryoprotected in Tissue-Tek optimal cooling temperature (OCT) compound for sectioning and cooled to a temperature of −20°C for sectioning. 40 μm coronal sections were cut using a cryostat. Brain sections were washed and permeabilized with PBST (0.4% Triton X-100 in PBS) over 15 min and incubated with blocking solution (5% normal goat serum in PBST) for 1 h at room temperature. Sections were incubated with rabbit α-mCherry or goat α-GFP polyclonal antibodies described in the key resources table overnight at 4°C. The following day slices were washed in PBST over 15 min and incubated in the AlexaFluor555-fluorescently-conjugated donkey α-rabbit secondary (1:500, Invitrogen) at RT for 1 h. Sections were mounted on slides with Vectashield Antifade Mounting Media with DAPI (Vector Laboratories). Slices were imaged using an Olympus Slideview VS200 slide scanning microscope.

### Behavioral Experiments

#### Behavioral hardware/software

Liquid reward tasks were conducted using MicroPython controlled behavioral boxes (pyControl^[27]^) with custom Python scripts written for each behavioral task. pyControl boxes contained 3 liquid delivery reward ports each with IR sensors for detecting port entry/exit, programmable LEDs for visual stimuli presentation, a programable speaker for presentation of auditory stimuli, and programmable solenoids for controlled liquid delivery. Functionality of each PyControl behavioral system was confirmed before each experimental trial. Data acquired from pyControls were analyzed using custom Python scripts to quantify reward port entries and reward port entry dynamics.

Fear conditioning was conducted in a Med Associates NIR Video Fear conditioning system and analyzed with the Video Freeze software from Med Associates. Researchers were blinded to the experimental conditions during behavioral analysis.

#### DREADD Agonist 21 (C21) administration

Mice were habituated to handling and scruffing for 3 days prior to initiation of DREADD agonist 21 delivery (C21, Tocris Biosciences, 0.3 mg/kg). Mice were fasted and water-restricted the night before experiments. Intraperitoneal injections of C21, were performed 15 minutes before initiation of behavioral conditioning sessions.

#### Cued-reward conditioning

Cued reward conditioning consisted of paired visual (LED within the reward port) and auditory cues predicting delivery of 10 µl of 10% sucrose. Trials were presented every 15 seconds over a 45-min session, yielding 72 cue–reward pairings. Reward-port entries were detected via infrared beam breaks. Entries occurring within 5 seconds following sucrose delivery were classified as cued reward responses.

#### Instrumental conditioning

Instrumental conditioning trials required mice to initiate each trial by nose-poking into the center reward port. Trial initiation triggered illumination of the left reward port, followed by delivery of 10 µl of liquid sucrose into the left reward port. Reward-port entries were detected via infrared beam breaks. Entries into the reward port within 5 seconds following initiation port entry were classified as instrumental responses.

#### Reward valuation

Reward valuation was structured such that either 10% sucrose flavored with 0.9 g/L of cherry flavored Kool-aid powder or 5% sucrose with 0.39 g/L of grape flavored Kool-aid powder were dissolved in tap water. On the first day of the experiment mice received IP saline injection (10 µl/g body weight) and were allowed access to both solutions. On days 2 and 3 mice received C21 (0.3 mg/kg) 15 minutes prior to the sessions and were only allowed access to the 5% grape flavored solution. On day 4 mice were again injected with saline prior to the session and allowed access to both solutions. Reward-port entries were detected via infrared beam breaks.

#### Reversal learning

Reversal learning experiments were similar to reward valuation experiments in that mice were provided access to both a cherry flavored 10% sucrose solution and a grape flavored 5% sucrose solution. However, in this experiment mice received C21 and access to both solutions throughout the experiment. After three days of conditioning the position of the rewards were switched and assessed for three more days. Reward-port entries were detected via infrared beam breaks.

### Statistics

Sample sizes for experiments were determined based on sample sizes used in similar experiments reported previously in the literature. Statistical analysis was conducted using GraphPad Prism software in which distribution of the data was analyzed to determine the appropriate statistical test to be applied. The statistical test used for each comparison is described in the figure legends corresponding to the specific figure. Data are presented as the mean ± SEM unless otherwise noted with p < 0.05 being the cut-off for a result to be considered significant. ‘‘n’’ corresponds to the number of individual mice or neurons analyzed as indicated.

## 3. Results

### 3.1. Cued reward conditioning increases inhibitory synaptic input onto Vglut1^+^ BLA→NAc neurons

To assess whether reward conditioning (Fig. 1A-D) alters synaptic inhibition of Vglut1^BLA→NAc^ neurons, we injected a Cre-dependent retrograde viral reporter (AAVrg-hSyn-DIO-eGFP) into the NAc of Vglut1-Cre mice to selectively label Vglut1^+^ neurons projecting to the NAc (Fig. 1E). Whole-cell recordings were performed from labeled neurons in the BLA in acute brain slices from naïve and reward–conditioned mice. Conditioned mice exhibited a significant increase in both the amplitude (Fig. 1G) and frequency (Fig. 1H) of spontaneous inhibitory postsynaptic currents (sIPSCs), consistent with enhanced inhibitory synaptic input onto Vglut1^BLA→NAc^ neurons. Additionally, resting membrane potential was significantly more hyperpolarized in conditioned neurons compared to naïve controls (Fig. 1I), suggesting a shift in intrinsic excitability. Although spontaneous firing rate showed a trend toward reduction (p = 0.0529; Fig. 1F), Vglut1^BLA→NAc^ neurons in reward conditioned mice displayed delayed spike initiation (Fig. 1J) and a reduced number of action potentials in response to current injection (Fig. 1K) relative to naïve mice, indicating a functional suppression of excitability of Vglut1^BLA→NAc^ neurons associated with reward conditioning. Together, these findings demonstrate that Vglut1^BLA→NAc^ neurons undergo inhibitory remodeling with sustained reward conditioning, characterized by increased synaptic inhibition, membrane hyperpolarization, and reduced excitability.

**Figure 1.**
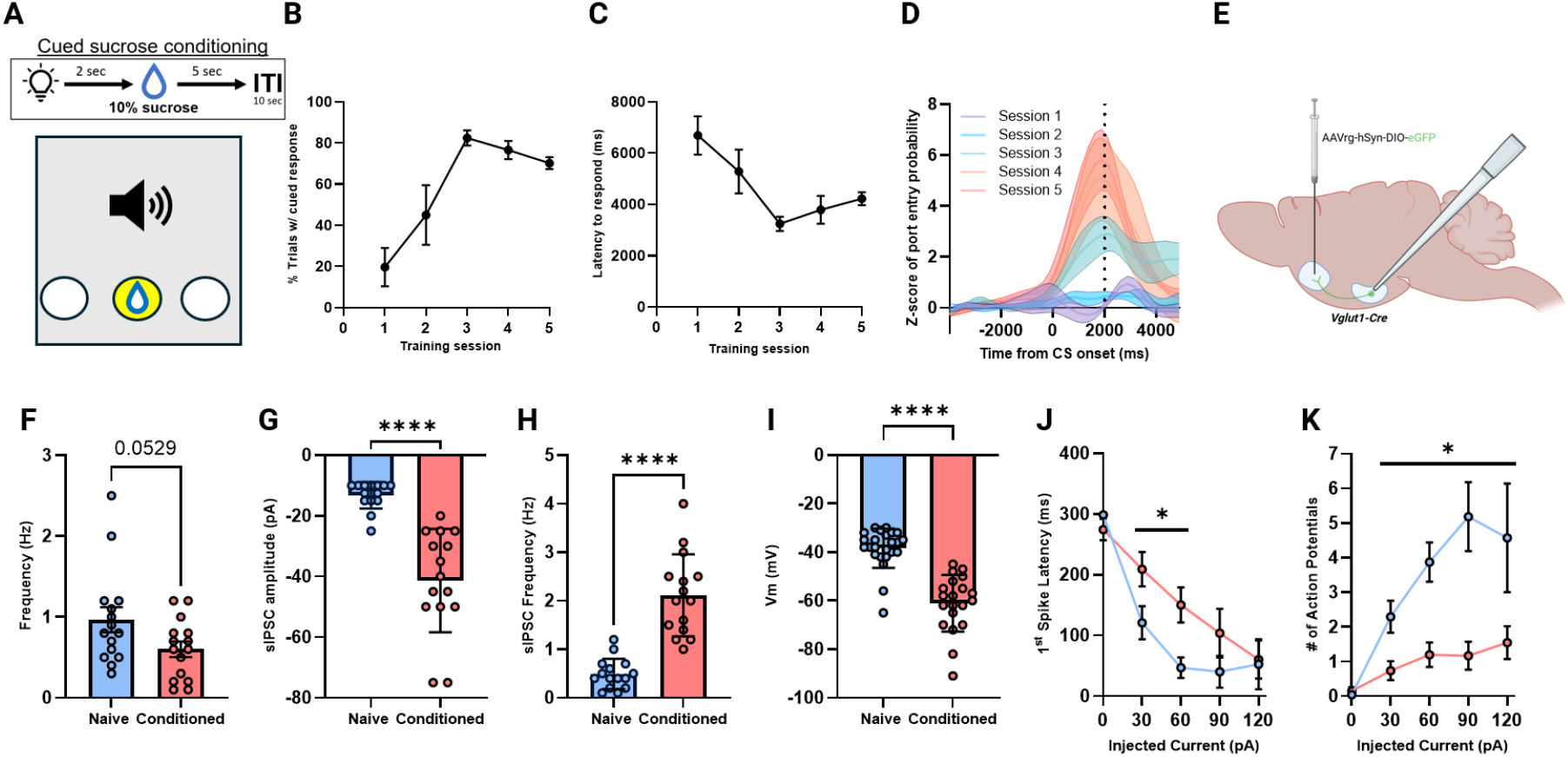
Reward conditioning increases inhibitory input onto Vglut1^BLA→NAc^ neurons. **(A)** Schematic of the cued sucrose conditioning task. A compound cue (tone and LED) predicts delivery of 10% sucrose to the center reward port 2 seconds after cue onset, followed by a 5 second consumption period, with an intertrial interval (ITI) of 10 seconds. **(B)** Average percentage of trials with a cued response (reward port entry within 7 seconds following cue presentation) across 5 training sessions (n = 6 mice). **(C)** Average latency to initiate a cued response across 5 training sessions from the same mice in B. **(D)** Average time-resolved z-scored probability of port entry aligned to cue onset across training sessions from the same mice in B. **(E)** Schematic of experimental design for fluorescently labeling BLA→NAc^Vglut1^ neurons using a retrograde AAV expressing Cre-dependent eGFP in Vglut1-Cre mice. **(F)** Average spontaneous action potential firing frequency of identified Vglut1^BLA→NAc^ neurons in naïve control mice and reward conditioned mice (n = 15 cells/group from 3 mice/group; *p* =0.0529, unpaired two-tailed t test). **(G)** Average spontaneous inhibitory post-synaptic current (sIPSC) amplitude in Vglut1^BLA→NAc^ neurons in naïve and conditioned mice (n = 15 cells/group from 3 mice/group; ****=*p* < 0.0001, unpaired two-tailed t test). **(H)** Average sIPSC frequency in Vglut1^BLA→NAc^ neurons in in naïve and conditioned mice (n = 15 cells/group from 3 mice/group; ****=*p* < 0.0001, unpaired two-tailed t test). **(I)** Average resting membrane potential (Vm) of Vglut1^BLA→NAc^ neurons in naïve and conditioned mice (n = 20-24 cells/group from 3 mice/group \*\*\*\**=p* < 0.0001, unpaired two-tailed t test). **(J)** Average first spike latency in response to current injection in Vglut1^BLA→NAc^ neurons in naïve and conditioned mice (n = 20-24 cells/group from 3 mice/group; *=*p* < 0.05, two-way ANOVA w/ Holm-Sidak’s multiple comparisons test). **(K)** Average number of evoked action potentials in response to current injection in Vglut1^BLA→NAc^ neurons in naïve and conditioned mice (n = 20-24 cells/group from 3 mice/group; *=*p* < 0.05, two-way ANOVA w/ Holm-Sidak’s multiple comparisons test).

### 3.2. Chemogenetic inhibition of Vglut1^BLA→NAc^ neurons enhances cued and instrumental reward responding

To investigate the behavioral consequences of inhibiting Vglut1^BLA→NAc^ neurons during reward conditioning, we used a dual-recombinase approach to enable specific chemogenetic inhibition of this population (Fig. 2A–D) during cued reward conditioning (Fig. 2E). Inhibition of Vglut1^BLA→NAc^ neurons markedly enhanced the acquisition of cue-evoked reward port entries and sustained this elevated performance across training sessions (Fig. 2F) relative to control mice (mice receiving C21 in the absence of AAVrg-DIO-Flp expression). Control mice gradually reached similar performance by later sessions, but inhibition of Vglut1^BLA→NAc^ neurons accelerated the development of cue-evoked responding and shifted the timing of port entries toward anticipatory responses as training progressed (Fig. 2G-I). To test whether this effect extended to action– outcome conditioning, mice were trained in an operant task requiring a nose poke to initiate reward delivery (Fig. 3A). Across training, inhibition of this projection significantly enhanced both initiation and reward port entries relative to controls (Fig. 3B,C), indicating increased task engagement. Reward port entry probability analyses revealed that inhibition strengthened the temporal association between initiation and reward consumption while also enhancing sustained reward-seeking across trials (Fig. 3D-F). Together, these findings indicate that suppression of Vglut1^BLA→NAc^ activity enhances both Pavlovian and instrumental components of reward conditioning, facilitating accelerated cue engagement and promoting sustained goal-directed actions.

**Figure 2.**
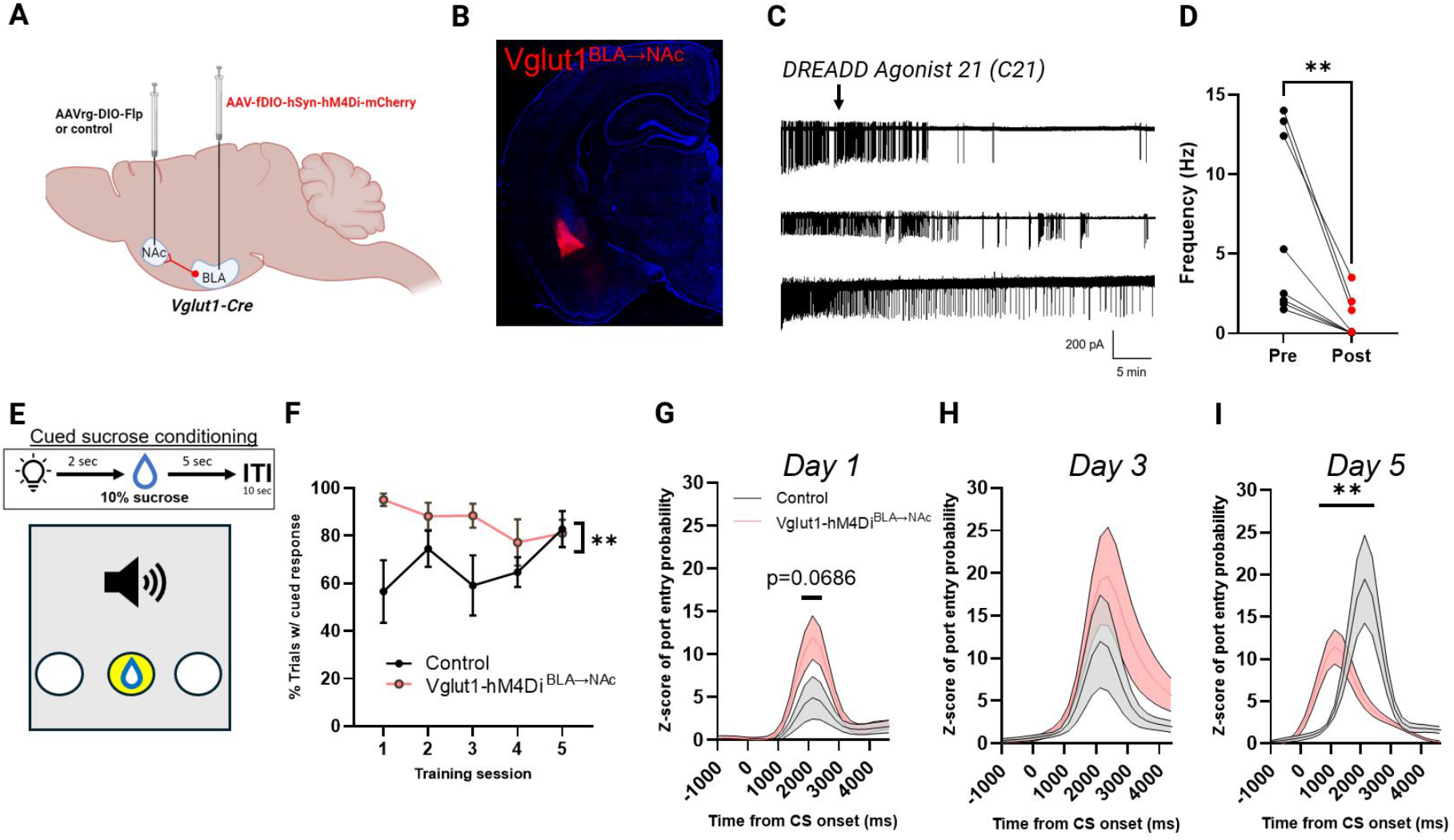
Chemogenetic inhibition of Vglut1^BLA→NAc^ neurons enhances cued sucrose responding and cue-evoked approach behavior. (A) Schematic illustrating the viral strategy used to selectively target Vglut1-expressing BLA→NAc projection neurons for chemogenetic inhibition. Vglut1-Cre mice received retrograde AAVrg-DIO-Flp (or control virus, AAVrg-DIO-EGFP) in the nucleus accumbens (NAc) and a Flp-dependent AAV-fDIO-hSyn-hM4Di-mCherry in the basolateral amygdala (BLA), restricting hM4Di expression to Vglut1^BLA→NAc^ projection neurons (Vglut1-hM4Di^BLA→NAc^). (B) Representative coronal brain section showing mCherry-labeled hM4Di expression in the BLA of Vglut1-Cre using the approach described in (A). (C) Representative whole-cell current-clamp recordings from hM4Di-expressing BLA neurons showing spontaneous firing before and after bath application of the DREADD agonist Compound 21 (C21). (D) Quantification of firing frequency before (Pre) and after (Post) C21 application in hM4Di-expressing BLA neurons (**p<0.01, paired one-tailed t test; n = 8 cells from 2 mice). (E) Schematic of the cued sucrose conditioning task. An auditory cue (2 seconds) predicted delivery of 10% sucrose (5 seconds), followed by an inter-trial interval (ITI; 10 seconds). (F) Percentage of trials with a cued response across five training sessions for control mice (black) and Vglut1-hM4Di^BLA→NAc^ mice (pink), (**p<0.01, two-way ANOVA with Holm–Sidak’s multiple comparisons; n = 8–11 mice/group). (G–I) Average time-resolved Z-scored probability of port entry aligned to cue onset for control (black/gray) and Vglut1-hM4Di^BLA→NAc^ (pink) mice on training day 1 (G), day 3 (H), and day 5 (I), (**p<0.01, two-way ANOVA with Holm–Sidak’s multiple comparisons). Shaded areas indicate mean ± SEM.

**Figure 3.**
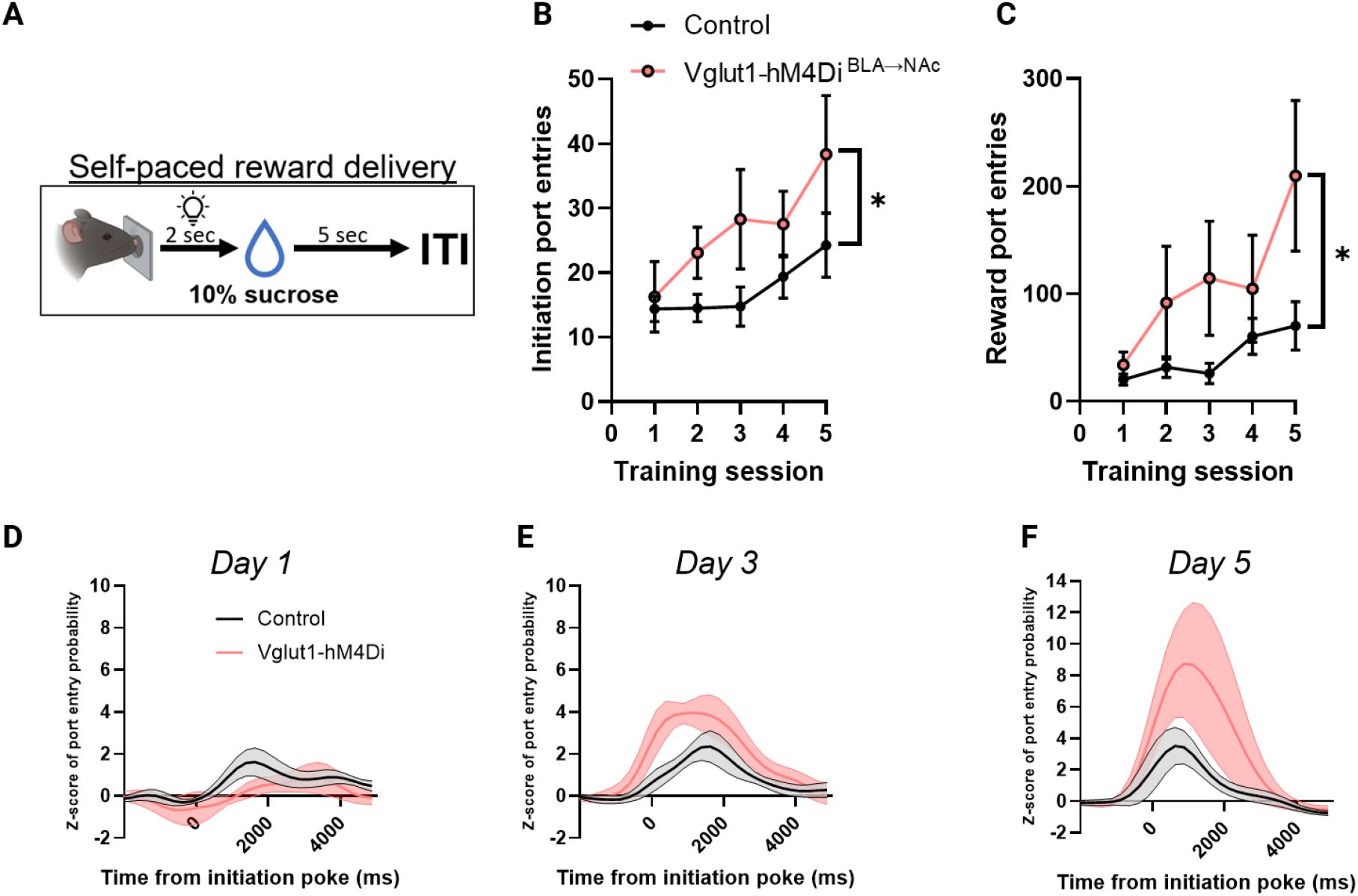
Chemogenetic inhibition of Vglut1^BLA→NAc^ neurons enhances instrumental reward responding. (A) Schematic illustrating the self-paced sucrose reward task. In the behavioral task, mice initiated trials via an initiation port, triggering a brief cue (2 seconds) followed by delivery of 10% sucrose (5 seconds) and an inter-trial interval (ITI, defined by next initiation port entry). (B) Mean number of initiation port entries across five training sessions for control mice (black) and Vglut1-hM4Di^BLA→NAc^ mice (pink), (*p<0.05, two-way ANOVA with Holm– Sidak’s multiple comparisons; n = 8–11 mice/group). (C) Mean number of reward port entries across training sessions from the same mice shown in B, (*p<0.05, two-way ANOVA with Holm–Sidak’s multiple comparisons). (D–F) Average time-resolved Z-scored probability of reward port entry aligned to the initiation poke for control (black) and Vglut1-hM4Di^BLA→NAc^ (pink) mice on training day 1 (D), day 3 (E), and day 5 (F). Shaded regions indicate mean ± SEM.

### 3.3. Inhibition of Vglut1^BLA→NAc^ neurons enhances reward port entry without altering discrimination or reversal learning performance

To distinguish whether inhibition of Vglut1^BLA→NAc^ neurons enhances reward-directed behavior through increased motivation or altered learning, we used a two-choice preference task in which mice received simultaneous cued access to cherry-flavored 10% sucrose and grape-flavored 5% sucrose delivered in separate ports with an acquisition and reversal phase. During acquisition, both experimental groups showed progressive increases in preference for the cherry-flavored 10% sucrose solution as expected (Fig. 4A). However, inhibition of Vglut1^BLA→NAc^ neurons increased combined reward port entries relative to control mice (Fig. 4B). During the reversal phase in which the positions of the rewards were switched both experimental groups adapted similarly (Fig. 4C), indicating preserved behavioral flexibility. This generalized elevation in reward port entries, coupled with intact acquisition and reversal performance, suggests that inhibition of Vglut1^BLA→NAc^ neurons increases overall reward-seeking activity, in concordance with our previous experiments, rather than altering discrimination or reversal learning processes.

**Figure 4.**
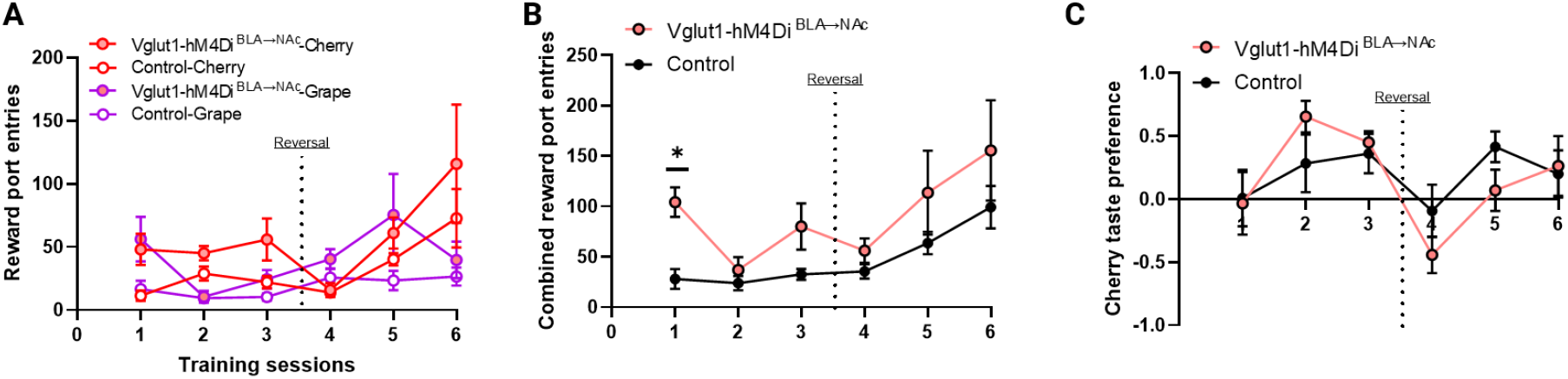
Chemogenetic inhibition of Vglut1^BLA→NAc^ neurons enhances reward port entry without altering discrimination or reversal learning performance. (A) Mean number of reward port entries for cherry-flavored 10% sucrose and grape-flavored 5% sucrose across training sessions in control and Vglut1-hM4Di^BLA→NAc^ mice. The dotted line indicates the reversal session, after which the location of the solutions were switched. (B) Combined reward port entries (collapsed across flavors) across training sessions for control (black) and Vglut1-hM4Di^BLA→NAc^ mice (pink), (*p<0.05, n=6-10 mice/group). (C) Cherry taste preference index across training sessions for control and Vglut1-hM4Di^BLA→NAc^ mice, calculated as the normalized difference in reward port entries between cherry- and grape-flavored solutions. Data are shown as mean ± SEM.

### 3.4. Inhibition of Vglut1^BLA→NAc^ neurons biases reward valuation toward a previously less-preferred option

To determine whether the motivational effects observed with inhibition of Vglut1^BLA→NAc^ neurons could reflect altered salience assignment, we used a two-choice taste preference task in which the initially less-preferred option was paired with chemogenetic inhibition (Fig. 5A). On day 1, mice were given simultaneous access to a cherry-flavored 10% sucrose solution and a grape-flavored 5% sucrose solution presented in separate ports in the absence of chemogenetic inhibition. On days 2–3, only the grape solution was available, and its consumption was paired with chemogenetic inhibition of Vglut1^BLA→NAc^ neurons. On day 4, mice were again given simultaneous access to both solutions in the absence of chemogenetic inhibition. As expected, both groups displayed a similar preference for cherry over grape on day 1 (Fig. 5B-F). On day 4, control mice maintained this preference for cherry, whereas mice in which Vglut1^BLA→NAc^ neurons were inhibited exhibited an increased preference for the grape flavored solution relative to control mice and on average displayed a similar preference for grape and cherry flavored solutions (Fig. 5G-K). These findings indicate that pairing inhibition of Vglut1^BLA→NAc^ neurons with the less-preferred grape solution attenuated the initial preference modifying the relative salience assigned to the two reward choices.

**Figure 5.**
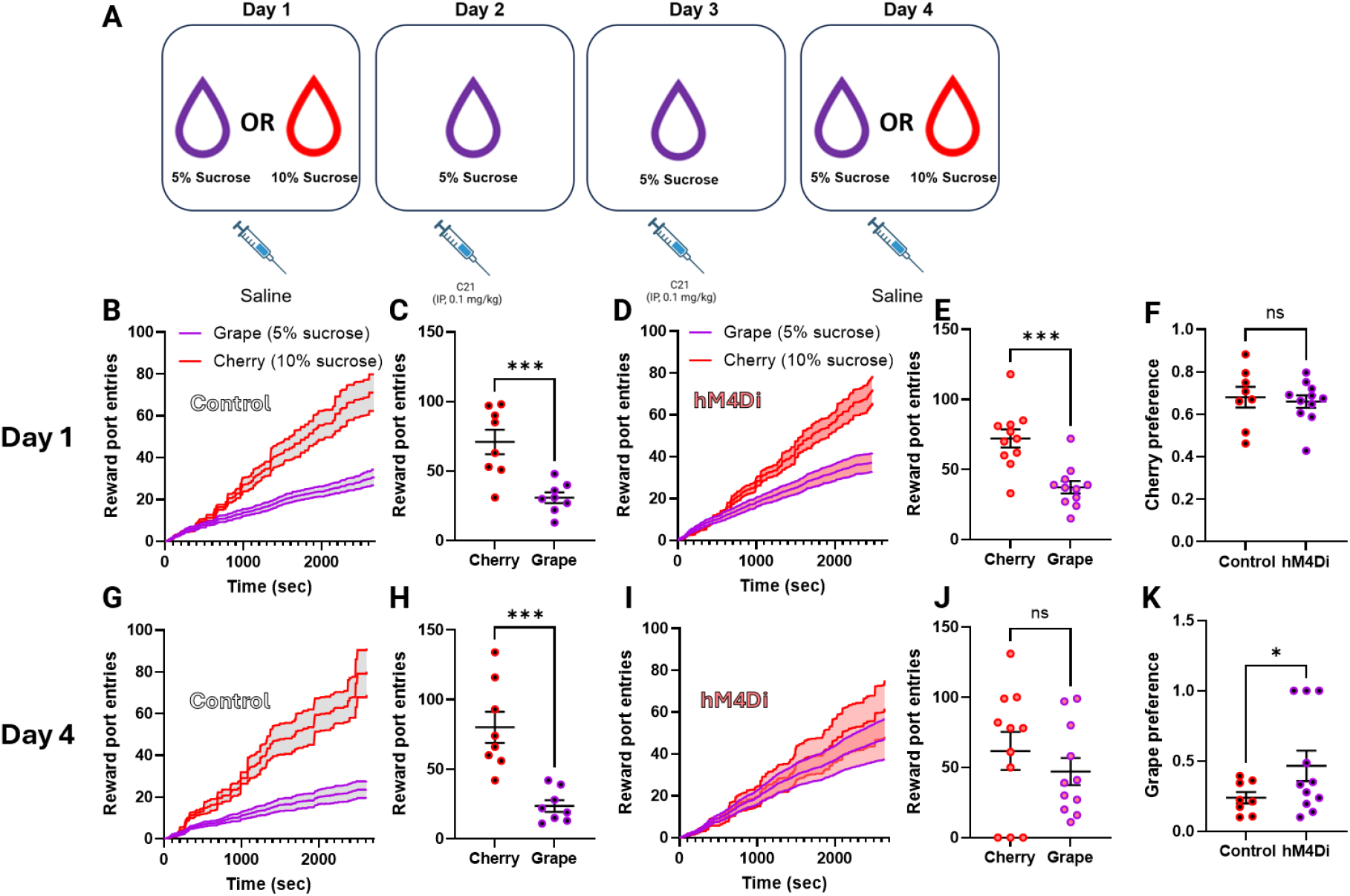
Chemogenetic inhibition of Vglut1^BLA→NAc^ neurons biases reward valuation toward a previously less-preferred option. (A) Schematic of the behavioral timeline. On Day 1, mice were given free access to two flavored sucrose solutions (5% sucrose with grape flavor vs. 10% sucrose with cherry flavor) to assess baseline preference following saline injection. On Days 2 and 3, mice received the DREADD agonist C21 (0.1 mg/kg, i.p.) prior to exposure to only the 5% sucrose grape solution. On Day 4, mice again received saline and were given free access to both sucrose solutions to assess preference following conditioning. (B–C) Average cumulative reward port entries over time (B) and total reward port entries (C) on Day 1 in saline-treated control mice (n = 8 mice; ***p < 0.001, unpaired one-tailed t test). (D–E) Average cumulative reward port entries (D) and total reward port entries (E) on Day 1 in Vglut1-hM4Di^BLA→NAc^ mice (n = 11 mice; ***p < 0.001, unpaired one-tailed t test). (F) Cherry preference index on Day 1 in control mice and Vglut1-hM4Di^BLA→NAc^ mice (ns, unpaired one-tailed t test). (G–H) Average cumulative reward port entries (G) and total reward port entries (H) on Day 4 in control mice (n = 8 mice; ***p < 0.001, unpaired one-tailed t test). (I–J) Average cumulative reward port entries (I) and total reward port entries (J) on Day 4 Vglut1-hM4Di^BLA→NAc^ mice (n = 11 mice; unpaired one-tailed t test). (K) Grape preference index on Day 4 in control mice and Vglut1-hM4Di^BLA→NAc^ mice (*p < 0.05). Individual data points represent individual mice. Data are shown as mean ± SEM.

### 3.5. Inhibition of Vglut1^BLA→NAc^ neurons enhances auditory fear discrimination

Given our results in the reward associated tasks suggesting inhibition of Vglut1^BLA→NAc^ neurons enhanced cued and instrumental reward acquisition likely through influencing salience assignment we wanted to determine whether these neurons exerted effects on salience assignment under aversive conditioning paradigms. To do so, we next examined inhibition of Vglut1^BLA→NAc^ neurons during discriminative cued fear conditioning. Briefly, during the conditioning day mice were exposed to two auditory cues one which predicted the delivery of a foot shock (CS^+^) and one which predicted the absence of a foot shock (CS^-^) (Fig. 6A). While inhibition of Vglut1^BLA→NAc^ neurons decreased overall freezing (Fig. 1B,C) we observed that the reduction in freezing was largely associated with the CS^-^ while no statistical difference was observed between the experimental groups when assessing the freezing in response to the CS^+^(Fig. 6D). Quantification of the animal’s ability to distinguish between the two cues (Discrimination Index, *((CS*^*+*^*-CS*^*--*^*)/(CS*^*+*^*+CS*^*-*^))) suggests inhibition of Vglut1^BLA→NAc^ neurons during fear conditioning enhanced discrimination between the CS^+^ and CS^−^ cues (Fig. 6E). Thus, inhibition of this specific subpopulation promotes more precise cue–outcome mapping across motivational contexts, suggesting a shared mechanism through which conditioning associated inhibition sharpens associative salience in both appetitive and aversive domains.

**Figure 6.**
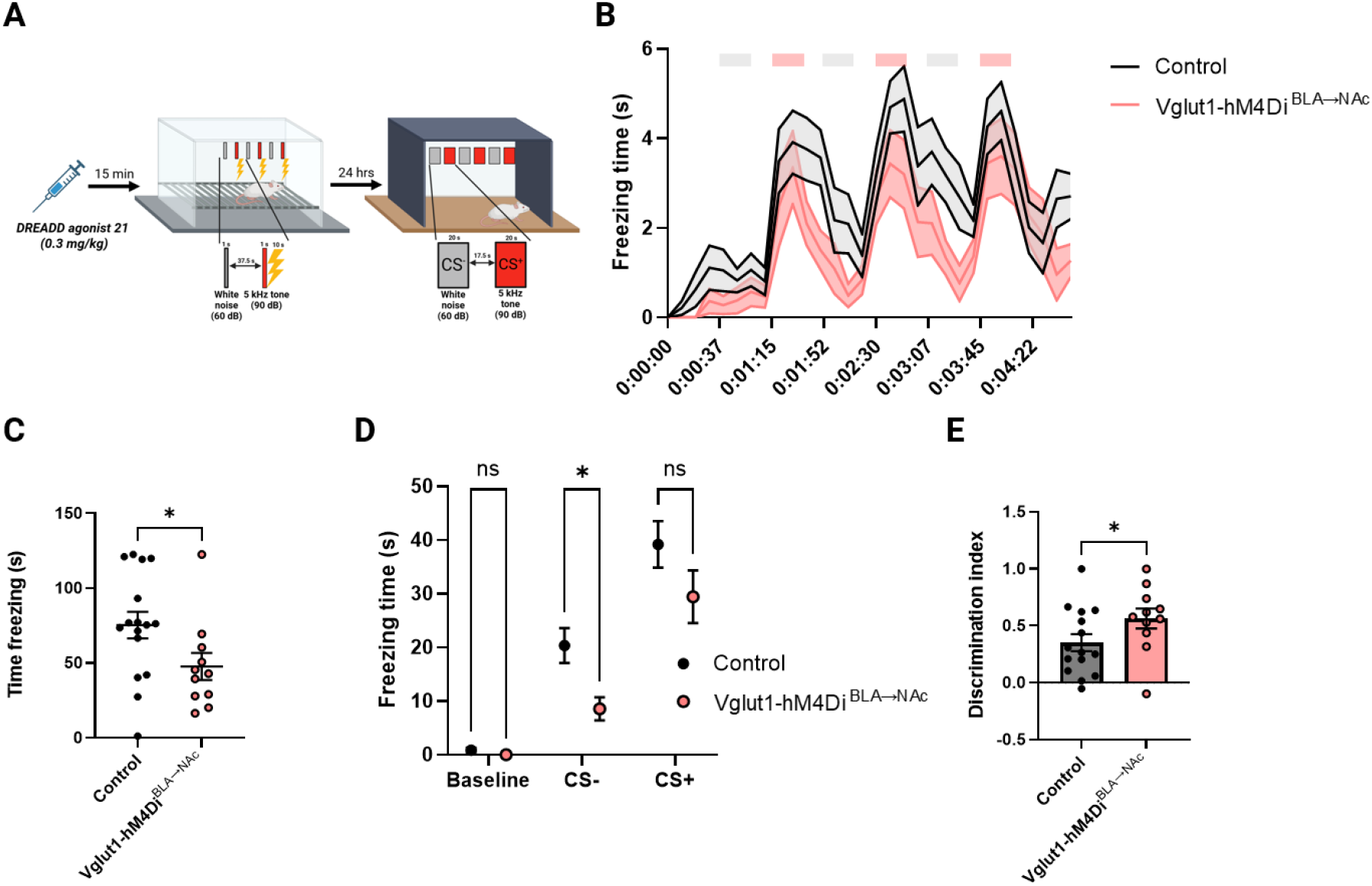
Chemogenetic inhibition of Vglut1^BLA→NAc^ neurons enhances auditory fear discrimination. (A) Schematic of the discriminative auditory fear conditioning and recall paradigm. Mice received the DREADD agonist Compound 21 (C21; 0.3 mg/kg, i.p.) 15 min prior to conditioning. During conditioning, mice were exposed to two auditory cues: a white noise (CS−, 60 dB) and a 5 kHz tone (CS+, 90 dB) paired with footshock. Twenty-four hours later, freezing behavior was assessed during presentation of CS− and CS+ in a distinct context. (B) Time-resolved freezing behavior during fear recall for control mice (black) and Vglut1-hM4Di^BLA→NAc^ mice (pink). Shaded bars indicate cue presentation epochs (grey = CS-, pink = CS+). (C) Total freezing time during the recall session collapsed across cue presentations, showing reduced freezing in mice with chemogenetic inhibition of Vglut1^BLA→NAc^ neurons relative to controls (*p < 0.05, unpaired two-tailed t-test). (D) Mean freezing time during baseline, CS−, and CS+ periods for control and Vglut1-hM4Di^BLA→NAc^ mice, (*p < 0.05, 2-way ANOVA w/ Dunnett’s multiple comparisons test). (E) Discrimination index [(CS+ − CS−)/(CS+ + CS−)] for control and hM4Di-BLA→NAc^Vglut1 mice, indicating enhanced discriminative fear expression in Vglut1-hM4Di^BLA→NAc^ mice (*p < 0.05, unpaired one-tailed t-test). Individual data points represent individual mice. Data are shown as mean ± SEM.

## 4. Discussion

Overall, our findings highlight a relatively underappreciated role for conditioning-associated inhibitory modulation of Vglut1^BLA→NAc^ neurons in shaping salience assignment to environmental stimuli during both appetitive and aversive conditioning. These results refine prevailing models of BLA→NAc function, which have largely emphasized how excitation of, or increased excitatory input onto, this projection facilitates appetitive and reward seeking behaviors. By demonstrating that inhibitory dynamics within a molecularly defined subset of BLA→NAc neurons are enhanced with conditioning and exert measurable behavioral consequences, our work adds mechanistic depth to the broader literature.

Supportive of our findings previous work has similarly observed that chemogenetic inhibition of BLA→NAc projecting neurons at-large enhances instrumental responding for sucrose reward^[28]^. Additionally, cholinergic suppression of BLA activity has also been reported to be sufficient to promote conditioned reward responding^[29]^ however, whether this is mediated through BLA→NAc projecting neurons or another projection defined population remains unresolved. However, there is a large body of previous literature suggesting excitation of BLA→NAc neurons promotes reward seeking behaviors which is somewhat at odds with our observations in the present work. It is important to note that within the BLA and NAc spatial subdivisions may play an important role in determining behavioral outcomes and therefore some of the apparent discrepancies may be due to differential targeting of BLA or NAc subregions which likely have diverse functional contributions to motivated behavior.

Regarding synaptic plasticity within the BLA during reward conditioning previous studies reported an increased AMPA/NMDA ratio in BLA→NAc neurons following reward conditioning, consistent with elevated excitatory drive^[7]^. However, those measurements were obtained after a single conditioning session and did not directly examine changes in intrinsic excitability or quantify inhibitory inputs, leaving open the question of how excitation-inhibition balance evolves as learning progresses. In contrast, our findings, obtained after multiple conditioning sessions, directly demonstrate enhanced increased inhibitory input and reduced excitability of Vglut1^BLA→NAc^ neurons. A potential synthesis of these observations is that conditioning promotes a temporal progression in synaptic regulation. Specifically, excitatory signaling may predominate early in conditioning driving appetitive behavior to support encoding of stimulus–outcome associations, whereas inhibitory signaling may emerge with continued conditioning to refine salience assignment, behavioral selection, and enhance precision as associations consolidate. In vivo single cell calcium imaging studies support conceptual framework demonstrating that as conditioning progresses the proportion of neurons in the BLA exhibiting inhibitory responses to stimuli predicting reward delivery increases^[23, 24]^.

However, in vivo electrophysiology specifically targeting BLA→NAc neurons following multiple conditioning sessions suggests a slightly different profile. Within BLA→NAc neurons a smaller proportion of neurons appear inhibited in response to reward predictive stimuli relative to those excited^[17]^. These results imply that increases in conditioning-associated inhibitory signaling observed in the BLA at-large may not be uniformly represented within BLA→NAc neurons. Our findings, which demonstrate a largely uniform increase in inhibitory regulation of Vglut1^BLA→NAc^ neurons, appears at odds with the prior projection-wide in vivo electrophysiology recordings. A key distinction is that our work specifically targeted Vglut1^BLA→NAc^ neurons as opposed to BLA→NAc neurons at-large. Moreover, we did not directly investigate cue-responsiveness, but rather general excitability of Vglut1^BLA→NAc^ neurons however, this will be an emphasis of our future studies. Potentially, both mechanisms are involved to further fine-tune behavioral responses.

Although Vglut1^BLA→NAc^ neurons appear to comprise the majority of glutamatergic BLA→NAc neurons^[25]^ a non-trivial percentage of BLA→NAc neurons are apparently GABAergic (∼30%^[25]^) with their role in motivated behaviors less well-defined. These observations strongly suggest the existence of functionally heterogenous subpopulations within BLA→NAc neurons with considered with additional evidence suggesting BLA→NAc projections defined by *Rspo2* and *Fezf2* expression contribute to defensive responses and aversive conditioning processes^[21, 22]^. Together, these findings underscore the need to move toward an integrated, molecularly and anatomically resolved framework for understanding how BLA circuits guide motivated behavior.

In summary, this work identifies Vglut1^BLA→NAc^ neurons as a molecularly distinct population whose inhibition modifies motivated behavior revealing an unexpected role for suppression within a classically excitatory projection. These results add to existing models of BLA→NAc function by demonstrating that inhibitory modulation within defined subpopulations fine-tunes motivated behavior likely through influencing salience assignment to environmental stimuli. From a translational perspective, these findings provide insight into mechanisms underlying maladaptive salience attribution in neuropsychiatric disorders^[30]^. Dysregulated inhibitory balance within the BLA→NAc circuit could disrupt salience assignment, as seen in anxiety and post-traumatic stress disorder, as well as addiction and obesity. By identifying an inhibitory mechanism that enhances associative precision and motivational focus, this work provides a framework for developing targeted interventions to modify salience processing in disorders of emotion and motivation.

**Supplementary Figure 1.**
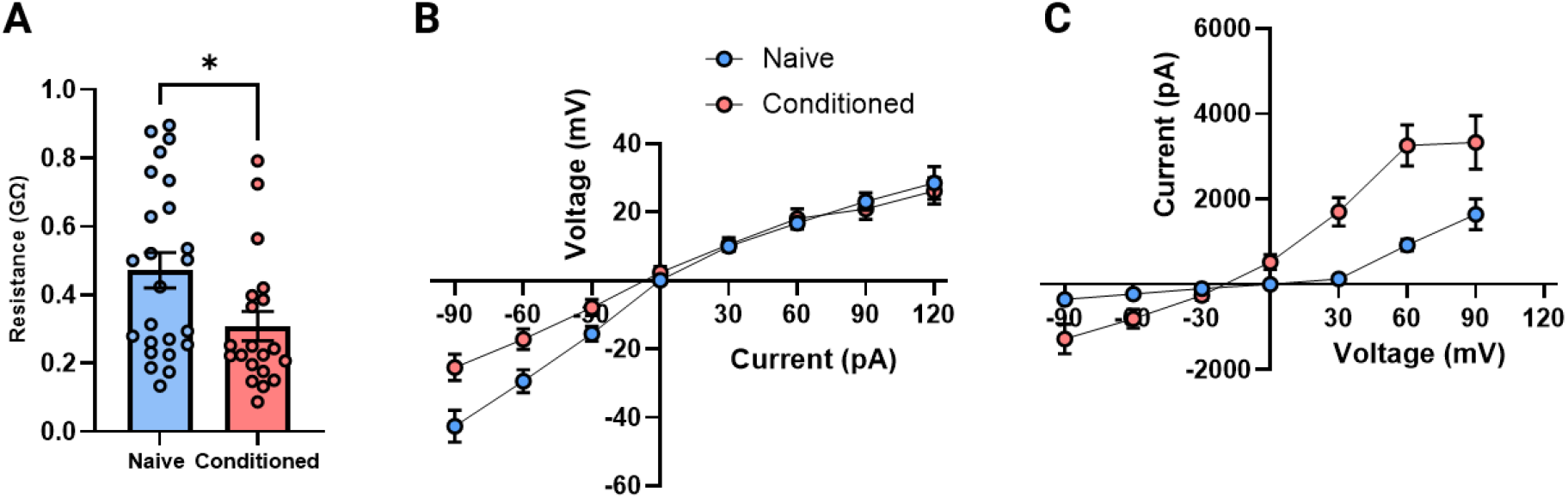
Reward conditioning is associated with altered intrinsic membrane properties of BLA→NAc neurons. (A) Input resistance measured in Vglut1^BLA→NAc^ projection neurons from naïve (blue) and reward-conditioned (pink) mice, (*p < 0.05, unpaired t test). (B) Current–voltage (I–V) relationship obtained under current-clamp conditions from Vglut1^BLA→NAc^ projection neurons in naïve and conditioned mice. (C) Voltage–current relationship measured under voltage-clamp conditions from the same populations of neurons. Each point represents an individual recorded neuron from 3 mice/group. Error bars indicate mean ± SEM.

## Bibliography & References Cited

1. Fanselow, M.S. and A.M. Poulos The neuroscience of mammalian associative learning. Annu Rev Psychol, 2005. 56, 207–34 DOI: 10.1146/annurev.psych.56.091103.070213.

2. Janak, P.H. and K.M. Tye From circuits to behaviour in the amygdala. Nature, 2015. 517, 284–92 DOI: 10.1038/nature14188.

3. Wassum, K.M. and A. Izquierdo The basolateral amygdala in reward learning and addiction. Neurosci Biobehav Rev, 2015. 57, 271–83 DOI: 10.1016/j.neubiorev.2015.08.017.

4. Romanski, L.M., et al. Somatosensory and auditory convergence in the lateral nucleus of the amygdala. Behav Neurosci, 1993. 107, 444–50 DOI: 10.1037//0735-7044.107.3.444.

5. Simmons, D.A. and D.B. Neill Functional interaction between the basolateral amygdala and the nucleus accumbens underlies incentive motivation for food reward on a fixed ratio schedule. Neuroscience, 2009. 159, 1264–73 DOI: 10.1016/j.neuroscience.2009.01.026.

6. Stuber, G.D., et al. Excitatory transmission from the amygdala to nucleus accumbens facilitates reward seeking. Nature, 2011. 475, 377–80 DOI: 10.1038/nature10194.

7. Namburi, P., et al. A circuit mechanism for differentiating positive and negative associations. Nature, 2015. 520, 675–8 DOI: 10.1038/nature14366.

8. Servonnet, A., P.P. Rompre, and A.N. Samaha Optogenetic activation of basolateral amygdala-to-nucleus accumbens core neurons promotes Pavlovian approach responses but not instrumental pursuit of reward cues. Behav Brain Res, 2023. 440, 114254 DOI: 10.1016/j.bbr.2022.114254.

9. Ambroggi, F., et al. Basolateral amygdala neurons facilitate reward-seeking behavior by exciting nucleus accumbens neurons. Neuron, 2008. 59, 648–61 DOI: 10.1016/j.neuron.2008.07.004.

10. Zhou, K., et al. Reward and aversion processing by input-defined parallel nucleus accumbens circuits in mice. Nat Commun, 2022. 13, 6244 DOI: 10.1038/s41467-022-33843-3.

11. Balleine, B.W., A.S. Killcross, and A. Dickinson The effect of lesions of the basolateral amygdala on instrumental conditioning. J Neurosci, 2003. 23, 666–75 DOI: 10.1523/JNEUROSCI.23-02-00666.2003.

12. Di Ciano, P. and B.J. Everitt Direct interactions between the basolateral amygdala and nucleus accumbens core underlie cocaine-seeking behavior by rats. J Neurosci, 2004. 24, 7167–73 DOI: 10.1523/JNEUROSCI.1581-04.2004.

13. Pitkanen, A., V. Savander, and J.E. LeDoux Organization of intra-amygdaloid circuitries in the rat: an emerging framework for understanding functions of the amygdala. Trends Neurosci, 1997. 20, 517–23 DOI: 10.1016/s0166-2236(97)01125-9.

14. Amorapanth, P., J.E. LeDoux, and K. Nader Different lateral amygdala outputs mediate reactions and actions elicited by a fear-arousing stimulus. Nat Neurosci, 2000. 3, 74–9 DOI: 10.1038/71145.

15. Nader, K., et al. Damage to the lateral and central, but not other, amygdaloid nuclei prevents the acquisition of auditory fear conditioning. Learn Mem, 2001. 8, 156–63 DOI: 10.1101/lm.38101.

16. Beyeler, A., et al. Organization of Valence-Encoding and Projection-Defined Neurons in the Basolateral Amygdala. Cell Rep, 2018. 22, 905–918 DOI: 10.1016/j.celrep.2017.12.097.

17. Beyeler, A., et al. Divergent Routing of Positive and Negative Information from the Amygdala during Memory Retrieval. Neuron, 2016. 90, 348–361 DOI: 10.1016/j.neuron.2016.03.004.

18. Blundell, P., G. Hall, and S. Killcross Lesions of the basolateral amygdala disrupt selective aspects of reinforcer representation in rats. J Neurosci, 2001. 21, 9018–26 DOI: 10.1523/JNEUROSCI.21-22-09018.2001.

19. O’Neill, P.K., F. Gore, and C.D. Salzman Basolateral amygdala circuitry in positive and negative valence. Curr Opin Neurobiol, 2018. 49, 175–183 DOI: 10.1016/j.conb.2018.02.012.

20. Paton, J.J., et al. The primate amygdala represents the positive and negative value of visual stimuli during learning. Nature, 2006. 439, 865–70 DOI: 10.1038/nature04490.

21. Kim, J., et al. Antagonistic negative and positive neurons of the basolateral amygdala. Nat Neurosci, 2016. 19, 1636–1646 DOI: 10.1038/nn.4414.

22. Zhang, X., et al. Genetically identified amygdala-striatal circuits for valence-specific behaviors. Nat Neurosci, 2021. 24, 1586–1600 DOI: 10.1038/s41593-021-00927-0.

23. Ottenheimer, D.J., et al. Orbitofrontal Cortex Mediates Sustained Basolateral Amygdala Encoding of Cued Reward-Seeking States. J Neurosci, 2024. 44, DOI: 10.1523/JNEUROSCI.0013-24.2024.

24. Zhang, X. and B. Li Population coding of valence in the basolateral amygdala. Nat Commun, 2018. 9, 5195 DOI: 10.1038/s41467-018-07679-9.

25. Jin, S., et al. Examining ventral subiculum and basolateral amygdala projections to the nucleus accumbens shell: Differential expression of VGLuT1, VGLuT2 and VGaT in the rat. Neurosci Lett, 2022. 788, 136858 DOI: 10.1016/j.neulet.2022.136858.

26. Ting, J.T., et al. Preparation of Acute Brain Slices Using an Optimized N-Methyl-D-glucamine Protective Recovery Method. J Vis Exp, 2018. DOI: 10.3791/53825.

27. Akam, T., et al. Open-source, Python-based, hardware and software for controlling behavioural neuroscience experiments. Elife, 2022. 11, DOI: 10.7554/eLife.67846.

28. Wang, Y., et al. A Critical Role of Basolateral Amygdala-to-Nucleus Accumbens Projection in Sleep Regulation of Reward Seeking. Biol Psychiatry, 2020. 87, 954–966 DOI: 10.1016/j.biopsych.2019.10.027.

29. Kimchi, E.Y., et al. Reward contingency gates selective cholinergic suppression of amygdala neurons. Elife, 2024. 12, DOI: 10.7554/eLife.89093.

30. Robinson, T.E. and K.C. Berridge The neural basis of drug craving: an incentive-sensitization theory of addiction. Brain Res Brain Res Rev, 1993. 18, 247–91 DOI: 10.1016/0165-0173(93)90013-p.

